# Segmented poly(A) tails with microRNA target sites confer tissue-specific regulation for mRNA therapeutics

**DOI:** 10.1101/2025.09.04.674363

**Authors:** Rui Qi, Rui Chen, Hua Chen, Ruiwen Xu, Lu Han, Yingmei Xu, Juan Li, Na Li, Qiang Li, Hui Bao, Tingting Zhang, Kai Lv, Yijie Dong, Shan Cen, Weiguo Zhang

**Affiliations:** RinuaGene Biotechnology Co., Ltd., Suzhou, Jiangsu Province, P. R. China; Institute of Medicinal Biotechnology, Chinese Academy of Medical Sciences and Peking Union Medical College, Beijing 100050, China

**Keywords:** poly(A), microRNA, miRNA target site, mRNA-LNP, selective expression, mRNA therapeutics

## Abstract

Targeted delivery and controlled expression of mRNA-LNPs are critical for the development of safe and effective mRNA medicines. However, efficient post-delivery regulation of mRNA-LNP expression remains challenging. In this study, we engineered segmented poly(A) tail variants that function as gene-specific regulatory elements for synthetic mRNAs. Specifically, we inserted microRNA target sites (MTS) for miR-122 or miR-142 at various positions of the poly(A) tail of synthesized luciferase reporter mRNA. These modifications significantly reduced luciferase expression in non-target tissues *in vitro* and *in vivo*, demonstrating position-dependent selective expression control. Furthermore, by incorporating triple-MTS sequences for miR-142, miR-126, and miR-148a in all possible combinations at the 5’ end of the poly(A) tail, we identified triple-MTS arrangements that simultaneously decrease luciferase activities in three non-target hepatic cell types, while preserving robust expression in hepatocytes. The results highlighted the importance of MTS insertion order for optimal mRNA silencing. These triple-MTS modules significantly expanded the utility of single miRNA-responsive elements, enabling cell-type selective regulation for mRNA therapeutics. By complementing tissue-tropic delivery LNPs, our mRNA cargo regulatory elements have the potential to improve tissue and cell type selectivity as a novel platform for post-delivery regulation of mRNA-LNP.

## Introduction

Messenger RNA (mRNA)-based technology demonstrated the huge potential as a platform for vaccines, protein replacement, and gene editing. The success of SARS-CoV-2 mRNA vaccines by Moderna and Pfizer/BioNTech highlights the utility of antigen-encoding mRNA encapsulated by lipid nanoparticles (LNPs) to elicit robust immune responses (*1, 2*). Beyond vaccines, mRNA-LNP technology has also been actively explored for treating rare genetic disorders and other diseases by delivering mRNAs encoding therapeutic proteins or gene-editing machinery (*3, 4*).

However, precise spatiotemporal control of mRNA expression remains a critical unresolved problem (*5*). Off-target expression can contribute to immune reactogenicity in vaccines (e.g., leaky expression of SARS-CoV-2 Spike protein in liver after intramuscular administration of mRNA vaccine) or toxicity in gene therapies (e.g., ectopic expression of gene editing tool in off-target tissues or cell types within the correct tissue) (*6*). Although advances in lipid discovery and LNP formulation facilitated the successful development of mRNA vaccines, achieving extrahepatic delivery or cell-type-specific control within hepatic tissues has been challenging due to an incomplete understanding of LNP biodistribution and cellular uptake mechanisms for different organs and tissues (*7*). Thus, novel strategies are needed to overcome the problem of off-target mRNA expression.

To address this limitation, a promising complementary strategy is to engineer the mRNA cargos to enhance tissue specificity (*8*). Synthetic mRNAs typically consist of an open reading frame (ORF) encoding the protein of interest and several essential regulatory elements, including 5′ cap, 5′ untranslated region (UTR), 3′ UTR, and a poly(A) tail for optimized expression (*9*). Particularly, the 3′ UTR naturally contains microRNA (miRNA) target sites (MTS) and other regulatory motifs that control mRNA stability and translation (*10*). miRNAs are a class of small non-coding RNAs showing tissue- and cell type-specific patterns (*11, 12*). After being incorporated into the RNA-induced silencing complex (RISC), they can mediate mRNA degradation or translational repression by binding complementary sequences in target mRNAs (*13*).

The miRNA-based mechanism has been utilized for tissue-selective expression of therapeutic mRNAs (*14*). By incorporating MTS into the 3′ UTR, several groups have successfully reduced off-target expression in mice. For instance, miR-122 is abundant in hepatocytes but absent in hematopoietic cells (*15*), while miR-142 shows the opposite pattern (*16*). miR-122 or miR-142 MTS inserted in the 3’ UTR showed significant silencing effects on mRNA expression in liver and spleen, respectively (*17*). Others have placed MTS in the 5′ UTR or ORF to achieve similar effects (*14, 18*). Circular RNAs with miRNA-responsive internal ribosome entry sites (IRES) further demonstrated the potential of miRNA-mediated regulation (*19*). Collectively, these studies underscored the utility of endogenous miRNA networks for mRNA therapeutics.

While the poly(A) tail is traditionally viewed as a passive stabilizer, recent work suggested that engineered poly(A) variants could unlock new regulatory dimensions. In eukaryotes, the poly(A) tail protects mRNA from exonuclease degradation and recruits poly(A)-binding protein (PABP) to enhance translation initiation (*20*). Although natural poly(A) tails lack gene-specific functions, their length and sequence can be modulated to facilitate mRNA manufacturing and protein yield. For example, a 120 nt poly(A) tail maximizes protein expression compared to those of shorter lengths (*21*), while a segmented A30-70 poly(A) tail variant developed by BioNTech improves template DNA stability in bacteria for IVT mRNA production (*22*). Cytosine-rich spacers or branched poly(A) structures have been shown to boost mRNA stability and expression (Li et al., 2022; Chen et al., 2025). However, these poly(A) engineering efforts primarily focus on preserving or enhancing its canonical functions for mRNA stabilization and translational regulation. Therefore, tissue-specific expression regulation through engineered poly(A) tail represents an excellent opportunity for expanding the regulatory capacity of mRNA medicines.

In this study, we modified the poly(A) tail into a novel miRNA-responsive regulatory element. We designed segmented poly(A) tail variants with embedded miR-122 or miR-142 target sites. These poly(A) variants potently suppressed mRNA expression in non-target tissues with efficacies impacted by their sites of insertion, while maintaining robust activity in desired organs *in vivo*. Furthermore, by sequentially incorporating multiple distinct MTS at the 5’ end of poly(A), we achieved simultaneous silencing effects in multiple hepatic cell types while allowing robust expression in cultured primary hepatocytes. This modular and straightforward platform holds promise for tissue-specific expression for safe mRNA medicines.

## Results

### Insertion of miRNA Target Sites in Poly(A) Tail Suppresses mRNA Expression in Cultured Cells

To systematically evaluate the silencing effects of microRNA (miRNA) target sites (MTS) at different 3’ terminal positions, we engineered firefly luciferase (*fLuc*) mRNAs containing either miR-122 or miR-142 MTS downstream of the α-globin 3’ UTR. These MTS were inserted at five distinct positions (0A, 14A, 19A, 30A, and 60A) within a segmented RG2 poly(A) variant, hereafter referred to as 122MTS and 142MTS, respectively (Fig. 1A). Control constructs included one that lacks any MTS sequences (hereafter referred to as noMTS) and two additional ones with either miR-122 or miR-142 MTS in the *α-globin* 3’ UTR (hereafter referred to as 122MTS-3UTR and 142MTS-3UTR, respectively) as previously reported (*17*).

**Figure 1.**
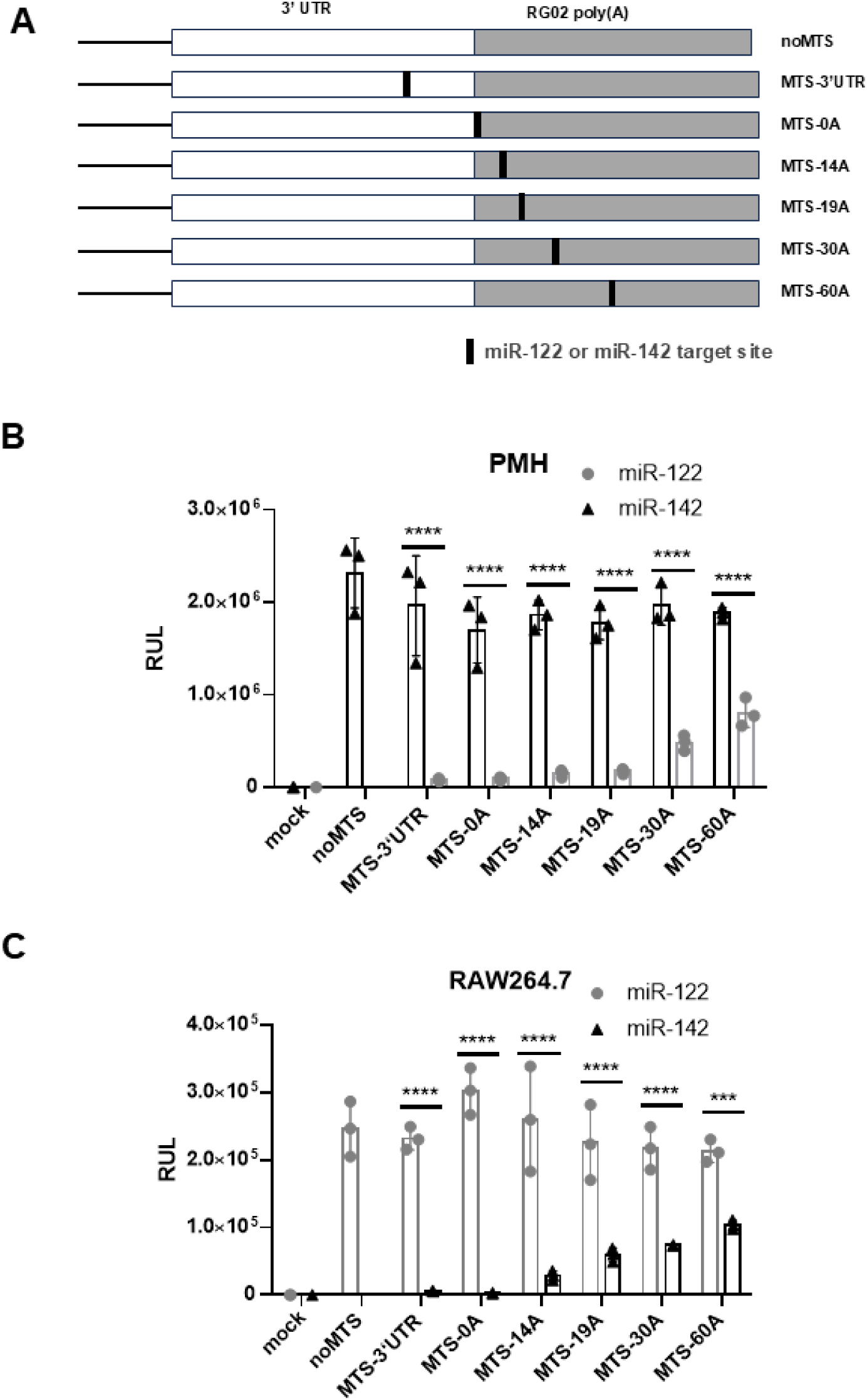
Single miR-122 and miR-142 target site insertions in the poly(A) tail of synthetic mRNAs specifically downregulate luciferase activity in non-target cell types, respectively. **(A)** Schematic diagram of miRNA target site (MTS) insertions for miR-122 and miR-142 at different positions of 3’ UTR and poly(A) downstream of the luciferase ORF. Only the 3’ UTR and the poly(A) regions were depicted. No insertion control (noMTS) and insertions in 3’ UTR (MTS-3UTR) or at five distinct positions of poly(A) were indicated as noMTS, MTS-3UTR, −0A, −14A, −19A, −30A, and −60A, respectively. **(B**) miR-122 target site insertions at distinct positions of the poly(A) tail effectively downregulate luciferase activity in primary mouse hepatocytes (PMH) compared to miR-142 target site insertions at the same locations or noMTS control. Bar graph shows the means of relative unit of luminescence (RLU) with SEM. **(C**) miR-142 target site insertions at distinct positions of the poly(A) tail effectively downregulate luciferase activity in RAW264.7 macrophages compared to miR-122 target site insertions at the same locations or noMTS control. Bar graph shows the mean of relative unit of luminescence (RLU) with SEM. *** p < 0.001; **** *p* < 0.0001 (one-way ANOVA).

We determined the impact of a single miR-122 or miR-142 target site inserted in the poly(A) tail on luciferase activity using primary mouse hepatocytes (PMHs, high miR-122 and low miR-142) and RAW264.7 macrophages (low miR-122 and high miR-142), based on their reciprocal expression patterns for the two miRNAs (*15, 16*). In PMH cells, all *fLuc*-142MTS constructs exhibited luciferase activities comparable to the *fLuc*-noMTS control, reflecting the low endogenous miR-142 levels in these cells (Fig. 1B). In contrast, all *fLuc*-122MTS constructs showed significant reductions in luciferase activity, consistent with miR-122-mediated silencing (Figs. 1B, S1). In addition, proximal insertions (0A, 14A, 19A) mediated strong silencing comparable to 122MTS-3UTR (*p* > 0.05, Student’s t-test), whereas distal insertions (30A, 60A) exhibited weaker suppression (Fig. 1B).

In RAW264.7 macrophages (high miR-142 and low miR-122), all *fLuc*-122MTS constructs displayed luciferase activities comparable to the noMTS control, consistent with the low endogenous miR-122 levels in these cells (Fig. 1C). Conversely, all *fLuc*-142MTS constructs showed significant reductions in luciferase activity with a position effect, mirroring the pattern observed for *fLuc*-122MTS in PMH (Fig. 1B). Two proximal insertions (142MTS-0A and −14A) mediated approximately 90% suppression in luciferase activity, comparable to 142MTS-3UTR (Figs. 1C, S1). In comparison, the three more distal insertions (19A, 30A, and 60A) exhibited progressively weaker silencing. Time-course RT-qPCR confirmed rapid degradation of *fLuc*-142MTS-0A mRNA in macrophages (Fig. S1). Together, these results demonstrate that MTS embedded in segmented poly(A) tails enable cell type-specific mRNA down-regulation, with silencing efficiency affected by both miRNA identity and insertion position.

### Combinatorial miRNA Targeting Inhibits Expression in Multiple Hepatic Cell Types while Preserving Expression in Hepatocytes

To minimize off-target mRNA expression in multiple cell types simultaneously, we engineered mRNAs with tandem miRNA target sites at the 0A position of the poly(A) tail, where single MTS insertions demonstrated strong, consistent silencing (Fig. 1). We designed two dual-MTS constructs containing both miR-122 and miR-142 target sites at the 0A position in different orders (122/142MTS-0A and 142/122MTS-0A; Fig. 2A). In both PMH cells and RAW264.7 macrophages, these constructs reduced luciferase activity to near-background levels, while the noMTS control maintained robust expression (Figs. 2B, 2C). These results demonstrate that the 3’ UTR-poly(A) junction can accommodate multiple MTS for potent, simultaneous silencing in different cell types, regardless of insertion order.

**Figure 2.**
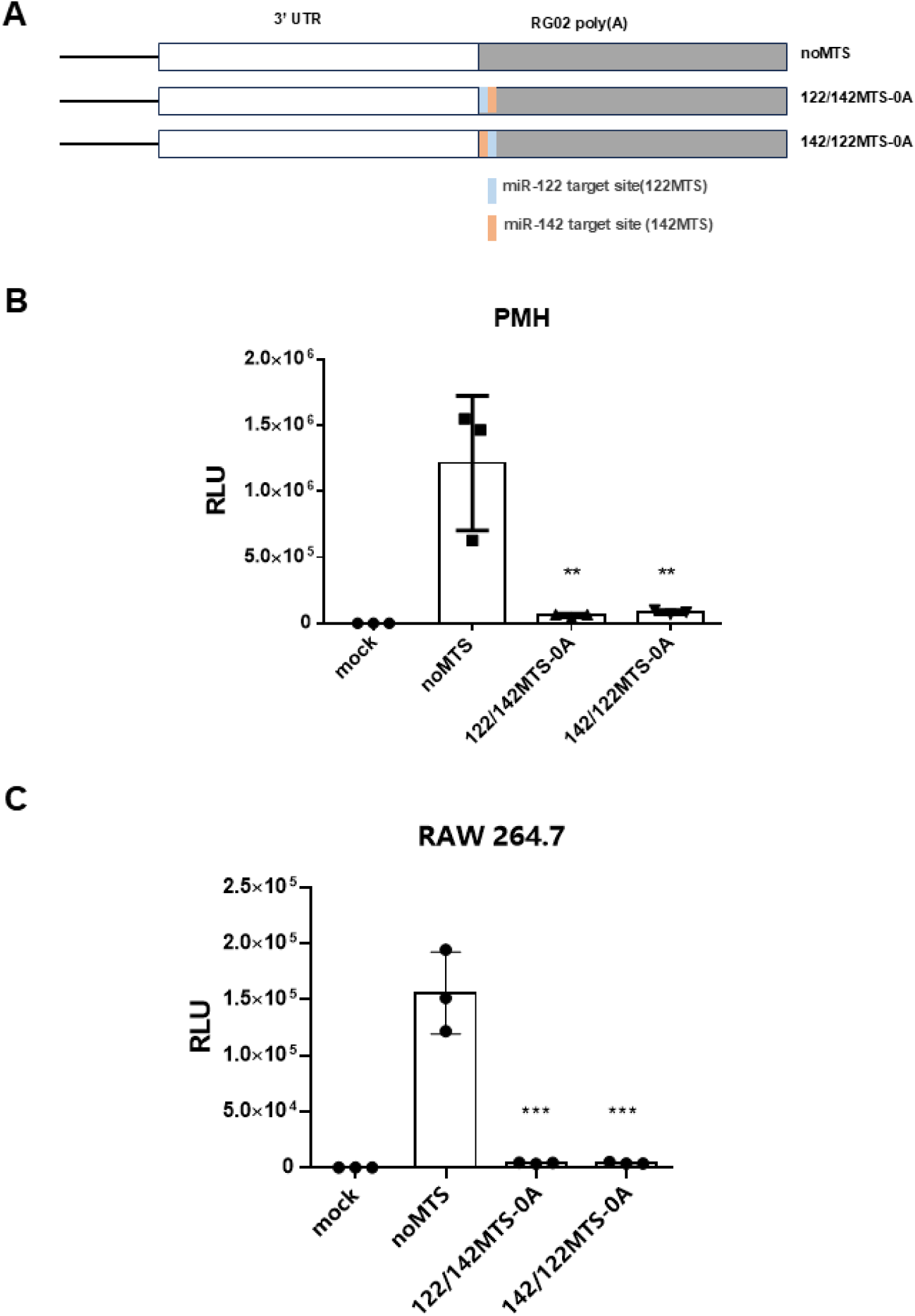
Dual-miRNA target sites for miR-122 and miR-142 at the 5’ end of poly(A) tail simultaneously downregulate mRNA expression in non-target cells. **(A**) Schematic diagram of dual miRNA target site (dual-MTS) insertions for miR-122 (blue) and miR-142 (orange) at 0A of position poly(A). Only the 3’ UTR and the poly(A) regions were depicted. No insertion control (noMTS) and the two insertions at 0A of poly(A) with opposite orders were indicated as noMTS, 122/142MTS-0A and 142/122MTS-0A. **(B-C**) Sequential insertions of miR-122 and miR-142 dual-MTS at 0A position of the poly(A) tail (122/142MTS-0A and 142/122MTS-0A) effectively downregulate luciferase activity in (**B**) primary mouse hepatocytes (PMH) and (**C**) RAW264.7 macrophages compared to noMTS control regardless of the order of insertion. Bar graph shows the means of relative unit of luminescence (RLU) with SEM (** *p* < 0.01; *** *p* < 0.001, one-way ANOVA).

To achieve silencing across multiple non-parenchymal liver cell populations while maintaining mRNA expression in hepatocytes, we engineered a triple MTS system targeting macrophages (miR-142), hepatic stellate cells (HSC) (miR-148a), and liver sinusoidal endothelial cells (LSEC) (miR-126) for silencing. We constructed all six possible MTS permutations at 0A position of the poly(A) tail (Fig. 3A). In PMH cells, which express low levels of all three miRNAs, five out of six triple-MTS configurations maintained luciferase activity comparable to the noMTS control as expected (*p* > 0.05, Student’s t-test; Fig. 3B). However, the 148a/142/126MTS-0A variant showed significantly reduced activity (59.0% suppression, *p* < 0.001, Student’s t-test), indicating that MTS order critically influences mRNA expression (Fig. 3B).

**Figure 3.**
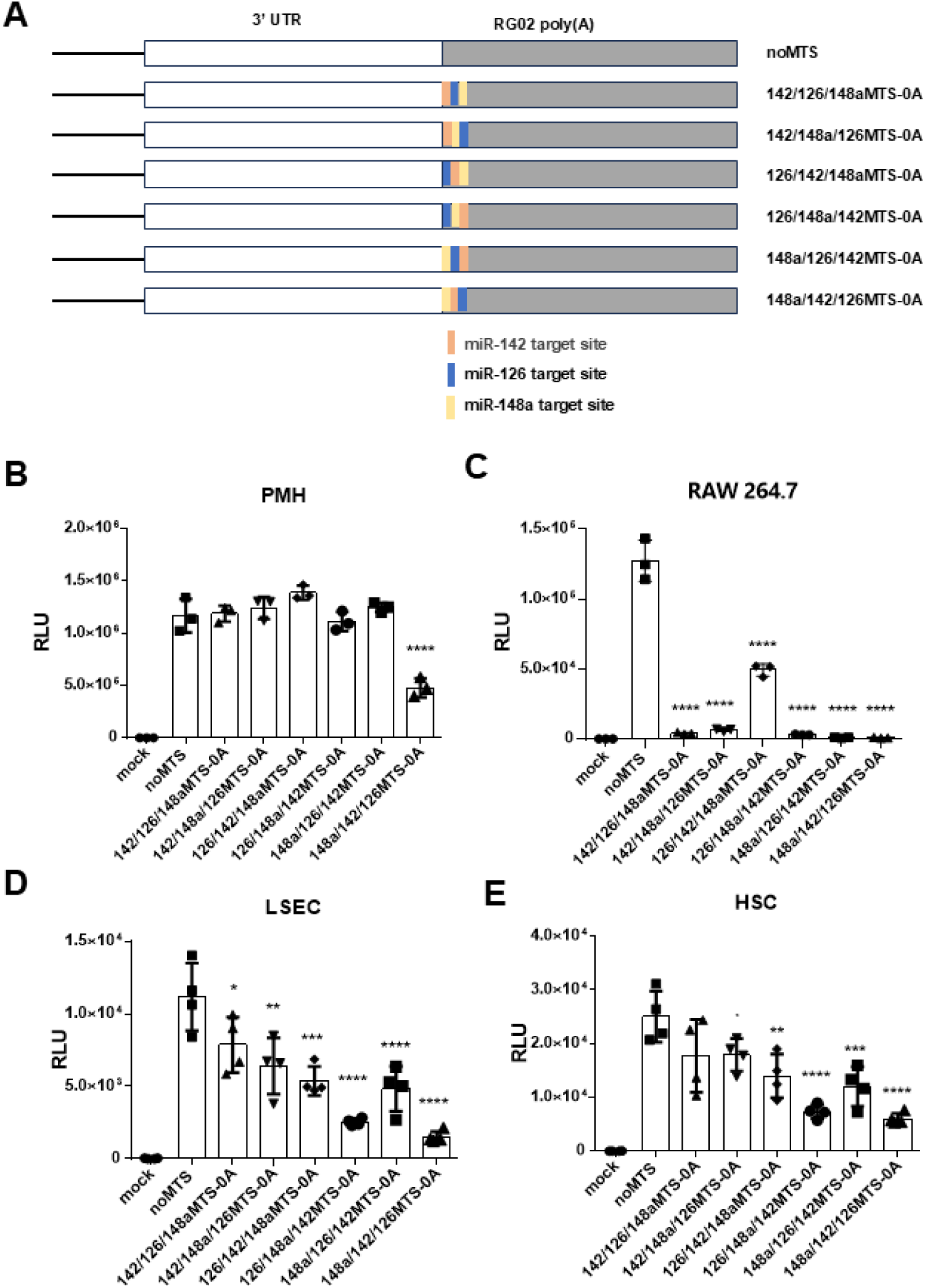
Triple miRNA target site (triple-MTS) insertions at the 5’ end of poly(A) tail allow effective expression in hepatocytes while simultaneously suppressing expression in multiple non-target hepatic cell types. **(A)** Schematic diagram of triple miRNA target site (triple-MTS) sequential insertions for miR-142 (orange), miR-126 (blue) and miR-148a (yellow) at 0A of position poly(A) with all possible permutations. Only the 3’ UTR and the poly(A) regions were depicted. No insertion control (noMTS) and all six possible permutations were indicated. **(B-E**) Bar graphs showing luciferase activity (RLU) for the negative control (mock), *fLuc* mRNA with no MTS control, and with triple-MTS at 0A position of the poly(A) tail in primary mouse hepatocytes (PMHs), RAW264.7, LSEC, and HSC, respectively. (**B**) In PMH cells, five out of six triple-MTS insertions effectively express luciferase at levels comparable to that of noMTS control with an exception showing reduction with extreme significance. (**C**) In RAW264.7 cells, all triple-MTS insertions show significant reductions in luciferase activity compared to the noMTS control, with five out of six showing more than 90% reductions. (**D**) In LSEC, all triple-MTS insertions displayed significant silencing compared to noMTS control, with one showing more than 70% reduction. (**E**) In HSC, five out of six displayed significant silencing compared to noMTS control, with one showing more than 70% reduction. Data were presented as mean ±SEM (* *p* < 0.05, ** *p* < 0.01, *** *p* < 0.001, **** *p* < 0.0001; one-way ANOVA).

In RAW264.7 macrophages, all triple-MTS combinations significantly reduced luciferase activity (*p* < 0.001, Student’s t-test), with five out of six configurations decreasing expression to <5% of the noMTS control (Fig. 3C). As an exception, 126/142/148aMTS-0A retained ∼33% activity. Similar patterns were observed in LSECs, where all constructs mediated significant suppression, with 126/148a/142MTS-0A showing maximal reduction (76% suppression; Fig. 3D). In HSCs, five configurations significantly decreased expression (69% maximal reduction for 126/148a/142MTS-0A), while 142/126/148aMTS-0A displayed insignificant reduction (Fig. 3E). Notably, 126/148a/142MTS-0A emerged as the overall most potent suppressor with less than 30% of the control activity for all three cell types (Figs. 3D, 3E).

Our comprehensive analysis demonstrates that most triple-MTS constructs effectively silence expression in macrophages, LSECs, and HSCs while preserving hepatocyte expression (Figs. S2). Among all tested configurations, 126/148a/142MTS-0A appeared to be the optimal arrangement, achieving simultaneous and robust suppression across all three non-parenchymal cell types. Notably, unlike the dual-MTS constructs, silencing efficiency in this triple-MTS system demonstrated strong dependency on MTS order, possibly reflecting configuration-specific mRNA secondary structures that influence miRNA accessibility.

### miRNA Target Sites Confer Poly(A) Tails Tissue-Specific Silencing Activity *in vivo*

To evaluate the *in vivo* silencing capacity of miRNA target sites embedded in poly(A), luciferase mRNAs containing miR-122 MTS at five distinct positions (0A, 14A, 19A, 30A, and 60A) of poly(A) were synthesized by IVT and encapsulated in (4S)-KEL12 lipid nanoparticles (LNPs). Six hours after intravenous (IV) administration, whole-body luminescence imaging revealed that the *fLuc*-noMTS control exhibited robust luciferase activity in the liver (Figs. 4A, 4B). (4S)-KEL12-based luciferase mRNA-LNP exhibits predominantly liver-specific luciferase activity when administered intravenously in our previous study (*23*). In contrast, the 3’ UTR-inserted 122MTS control (*fLuc*-122MTS-3UTR) showed 91.6% suppression compared to the noMTS control (*p* < 0.001, Student’s t-test), consistent with published data (*17*). Strikingly, all MTS configurations in poly(A) achieved significant silencing (*p* < 0.001), with efficacy strongly correlating with insertion proximity to the 3’ end (Figs 4A, 4B). The three most proximal insertions (0A, 14A, 19A) all mediated at least 90% suppression, with two surpassing even the 3’ UTR insertion control (95.8% for 0A, 94.7% for 14A, and 91.0% for 19A, respectively). In contrast, the distal 30A and 60A insertions showed progressively weaker inhibition (71.1% and 66.6%), suggesting that silencing potency decays as MTS sites are positioned deeper into the poly(A) tail.

**Figure 4.**
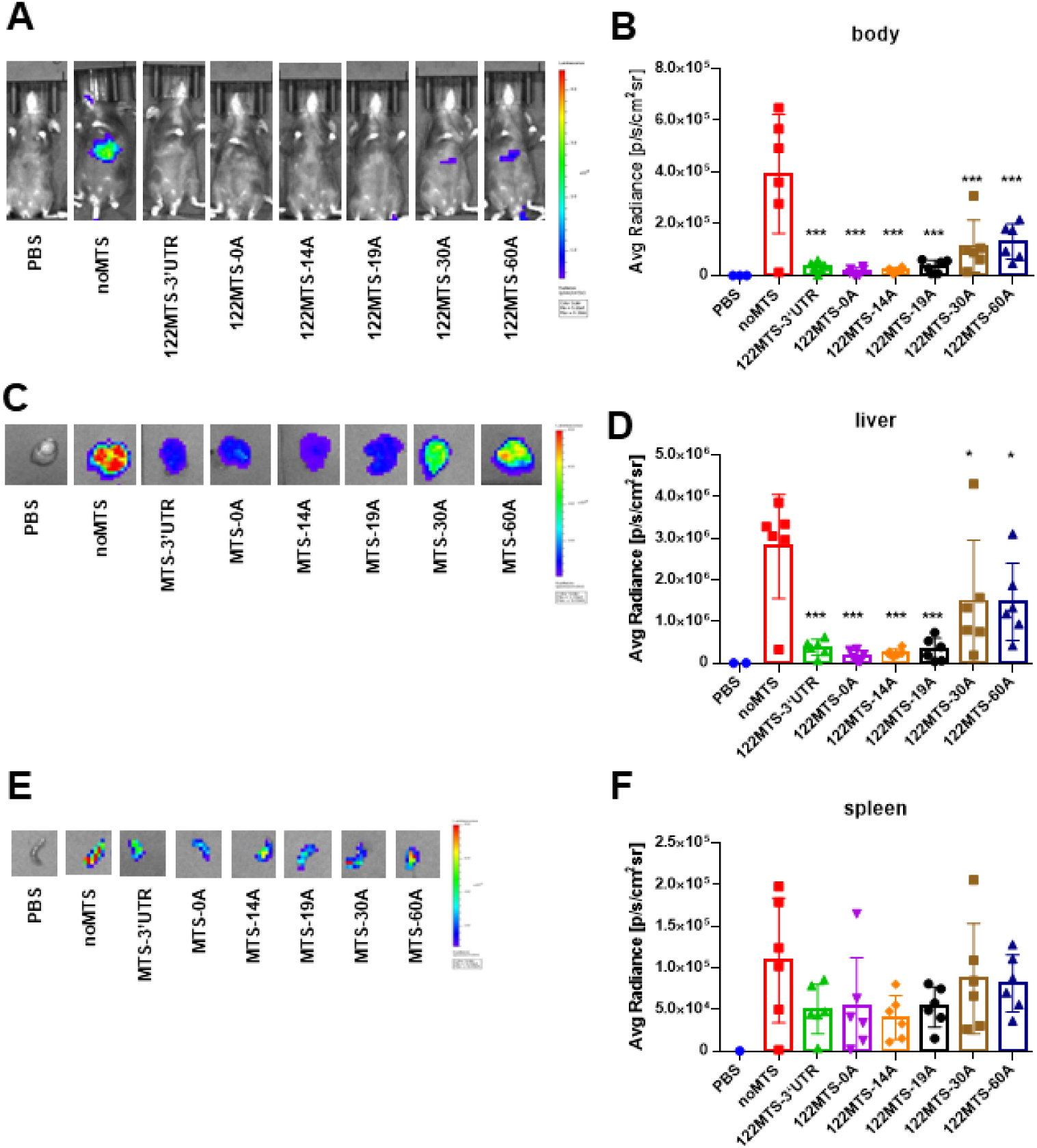
miR-122 target sites embedded in the poly(A) tail effectively suppress luciferase activity in mouse liver. Representative images of (**A**) whole-body, (**C**) livers, and (**E**) spleens from animals injected with KEL12-based LNPs encapsulating *fLuc* mRNAs containing no MTS insertion or miR-122 MTS inserted at distinct locations of the poly(A) tail six hours after tail vein injection (n=6). PBS was injected as negative control (n=3). (**B, D,F**) Bar graphs in are quantifications of average radiance of images in **A, C,** and **E**, respectively. Data were shown as mean, with the error bars representing SEM (* *p* < 0.05, ** *p* < 0.01, *** *p* < 0.001, **** *p* < 0.0001; one-way ANOVA).

Moreover, imaging using dissected organs unequivocally demonstrated that 122MTS embedded in poly(A) tail mediated strong silencing in liver (Figs. 4C, 4D). The proximal insertions (0A-19A) exhibited particularly potent suppression in liver luminescence (86.6-93.9% reductions, *p*<0.001, Student’s t-test). 122MTS-14A and - 19A achieved 93.9% and 91.1% silencing, respectively, surpassing the canonical 3’UTR MTS insertion control (86.6% silencing). This *in vivo* validation confirmed our findings in cultured cells and whole-body imaging experiments.

However, all *fLuc*-122MTS constructs exhibited measurable silencing in the spleen without statistical significance (20.2-53.6% reduction, *p* > 0.05, Student’s t-tests), suggesting some unexpected miR-122 activity in this organ (Figs. 4E, 4F). The observed decreases of expression in spleen according to insertion proximity (62.9% for 14A vs 25.3% for 60A) mirrored the trend in the liver, suggesting similar regulation with lower efficiency (Figs. 4E, 4F). Notably, spleen-to-liver signal ratios revealed that three proximal poly(A) insertions (0A-19A) achieved over 3-fold improved tissue selectivity compared to noMTS controls, matching or exceeding the canonical 3’ UTR MTS insertion control (Table 1). Therefore, we concluded that synthetic poly(A) tails with MTS insertions have the potential to functionally recapitulate and even surpass the regulatory capacity of 3’ UTRs.

**Table 1.** Relative activity of *luciferase* mRNA with 122MTS inserted at various locations downstream of 3’ UTR in dissected liver and spleen.

| *fLuc*-122MTS | Ratio (spleen/liver) | Normalized Ratio |
| --- | --- | --- |
| noMTS | 3.9% | 1.0 |
| 3' UTR | 13.4% | 3.5 |
| 0A | 31.0% | 8.0 |
| 14A | 16.1% | 4.2 |
| 19A | 16.4% | 4.2 |
| 30A | 5.8% | 1.5 |
| 60A | 5.5% | 1.4 |

We next studied whether miR-142 target sites engineered into the poly(A) tail could similarly confer tissue-specific regulation *in vivo*. Following IV delivery of KEL12 LNP-encapsulated *fLuc*-142MTS mRNAs, dissected organ imaging demonstrated potent spleen-specific silencing for all tested 142MTS constructs. Five of six 142MTS achieved 74% or more suppression in spleen (*p* < 0.001 vs noMTS control, Student’s t-test), with only the distal 142MTS-60A insertion showing much reduced silencing efficiency (47%) (Figs. 5A, 5B). Crucially, these 142MTS constructs maintained stable hepatic expression, as evidenced by both dissected liver imaging (Figs. 5C,5D) and whole-body luminescence (Figs. 5E,5F). Quantitative analysis revealed significantly enhanced liver-to-spleen expression ratios for all 142MTS constructs, confirming that poly(A)-embedded MTS can simultaneously preserve target liver tissue expression while suppressing off-target spleen activity (Table 2). Importantly, most poly(A)-inserted 142MTS displayed higher relative liver/spleen ratios than the canonical 142MTS-3UTR control. These findings collectively established poly(A) tail as a programmable platform for implementing tissue-specific gene regulation *in vivo*.

**Figure 5.**
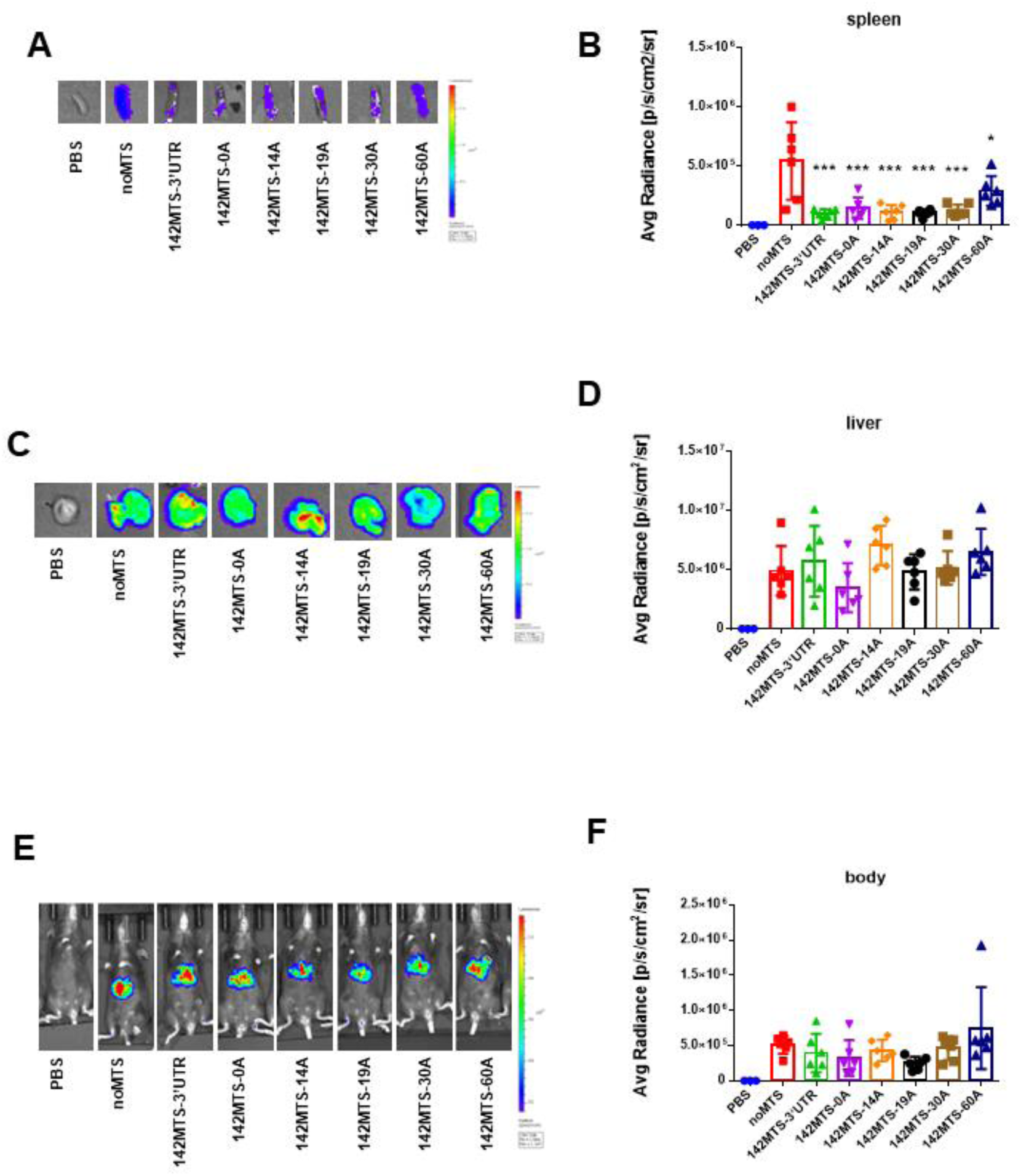
miR-142 target sites embedded in the poly(A) tail effectively suppress luciferase activity in mouse spleens. Representative images of (**A**) spleens, (**C**) livers, and (**E**) whole-body from animals injected with KEL12-based LNPs encapsulating *fLuc* mRNAs containing no MTS insertion or miR-142 MTS inserted at distinct locations of the poly(A) tail six hours after tail vein injection (n=6). PBS was injected as negative control (n =3). Bar graphs in **B, D,** and **F** are quantifications of average radiance of images in **A, C,** and **E**, respectively. Data were shown as mean, with error bars representing SEM (* *p* < 0.05, ** *p* < 0.01, *** *p* < 0.001, **** *p* < 0.0001; one-way ANOVA).

**Table 2.** Relative activity of *luciferase* mRNA with 142MTS inserted at various locations downstream of 3’ UTR in dissected liver and spleen.

| *fLuc*-142MTS | Ratio (Liver/Spleen) | Normalized Ratio |
| --- | --- | --- |
| noMTS | 9.0 | 1 |
| 3' UTR | 59.2 | 6.6 |
| 0A | 24.4 | 2.7 |
| 14A | 63.7 | 7.1 |
| 19A | 46.3 | 5.1 |
| 30A | 41.7 | 4.6 |
| 60A | 22.5 | 2.5 |

## Discussion

This study developed novel segmented poly(A) variants that can function as miRNA-responsive regulatory elements, expanding the toolbox for mRNA therapeutics. By embedding microRNA target sites within poly(A) tail, we modified poly(A) tail from a non-gene-specific element for mRNA stabilization and translation initiation to an active regulatory platform for tissue-specific gene silencing (Figs. 1, 4, and 5). We further explored a modular system using tandem MTS insertions at the 5’ end of poly(A) that simultaneously suppresses mRNA expression in multiple off-target liver cell types, such as macrophages, LSECs and HSCs, while maintaining hepatocyte expression *in vitro* (Figs. 2 and 3). Together, this work addresses a critical need for precise expression in mRNA therapies, particularly for applications requiring stringent control in off-target cell types, such as gene editing or immunogenic protein delivery (*24*).

Our insertion strategy within poly(A) has several potential advantages over previous methods of inserting MTS in the UTRs. Our approaches avoid disrupting the primary sequence of UTR. Although optimal MTS positioning in poly(A) requires experimental validation, our *in vitro* and *in vivo* results confirm silencing efficacies matching MTS inserted in the 3’ UTR while preserving translational capacity in desired organs (Figs. 4-5). This makes our approach an attractive alternative or even necessary for therapies where UTR integrity is critical (*25*). Finally, our approaches are unexclusive to MTS inserted in UTRs or LNPs with organ tropism such as SORT (*26*). Their integration could facilitate the development of more versatile organ- and cell type-selective expression vectors.

Our data suggested at least two critical factors affecting silencing efficiency for MTS inserted in poly(A). First, we observed stronger suppression with miR-122 compared to miR-142 (Fig. 1B vs 1C, and Fig. 4D vs 5B), which might reflect different abundances of these miRNAs in their respective tissues or cells. Second, positional analysis demonstrated more effective silencing for proximal MTS insertions than for more distal insertions in general (Figs. 1B-1C, 4D, and 5B). Although the exact mechanisms remain unclear, we envision a possible scenario where cleavage of poly(A) by the RISC complex recruited to proximal MTS in poly(A) likely results in shorter poly(A) tail length with less protective function than if recruited to a distal MTS, potentially responsible for better silencing efficacy for proximal insertions.

In addition, we observed some unexpected decreases of luciferase expression in the spleen using KEL12-based LNPs loaded with *fLuc* mRNAs carrying a miR-122 target site in the 3’ UTR or poly(A) tail. In contrast, miR-122 target site inserted at the same position in *α-globin* 3’ UTR did not show any reduction of luciferase activity in the spleen in an earlier study (*17*). We used a KEL12-based LNP that is different from the delivery system in Jain et al. It is possible that KEL-12 LNP might have a different cell type selectivity in the spleen from the earlier study.

Our *in vitro* studies also identified an optimal triple-MTS configuration (miR-126/142/148a) at the 5’ end of poly(A) that simultaneously suppresses mRNA expression in macrophages, LSECs and HSCs while maintaining robust hepatocyte expression (Figs. 2 and 3). Future validation of this triple-MTS *in vivo* will offer a convenient tool for hepatocyte-selective mRNA-LNP expression for protein replacement or gene editing. Although it remains unclear about order-dependent silencing efficacy for triple MTS insertions, it is possible that MTS arrangement may influence RISC accessibility through order-specific secondary structures.

Importantly, this modular platform offers opportunities for potential expansion to other tissue or cell type targets through rational MTS engineering. Together, the simplicity and modularity of our approaches including repurposed poly(A) tails and multiple MTS configurations suggest broad applications for cell-type-specific expression control, both within the liver and in extrahepatic tissues. Their future combinations with cell-specific delivery systems may unlock programmable expression control across diverse tissues.

## Materials and Methods

### Plasmids

All plasmids for in vitro transcription (IVT) used in this study were derived from a pUC57-Luc plasmid containing a T7 promoter region, 5’ α-globin UTR, firefly luciferase (fLuc) open reading frame (ORF), 3’ α-globin UTR, and a segmented poly(A) tail variant. The sequence of the poly(A) variant is as follows: 5’-AAAAAAAAAAAAAAAAAAAAAAAAAAAAAAAAAAAAAAAAAAAAAAAAAAAAAAAAAAAAGATATCAAAAAAAAAAAAAAAAAAAGAAAAAAAAAAAAAAAAAAAGAAAAAAAAAAAAAAAAA – 3’. The pUC57-Luc IVT template DNA plasmid containing the FLuc reporter gene was generated by gene synthesis service (Shanghai Dynegene Technologies Co., Ltd), using pUC57 as the vector backbone.

### Sequences of Modified Poly(A) Tail Variants

Sequences of the modified poly(A) variants were listed in Table S1.

### mRNA In Vitro Transcription (IVT) and Purification

Plasmid DNA was isolated from 50 mL of overnight bacterial culture using EndoFree Plasmid Maxi Kit (Qiagen, Cat. #12362), followed by BspQ1 (New England Biolabs, Cat.# R0712L) digestion to linearize DNA for IVT. IVT was performed using the High Yield T7 RNA Synthesis Kit (Hongene Biotech, Cat# ON-40) in a 100 µL reaction mixture of containing 5 µg of plasmid template, ATP, GTP, CTP, 1-N-Me-PseudoUTP (final concentration is 10 mM for each) and 5 µL of Cap analog (Synthgene, Cat.# CAP3011). The mixture was incubated at 37°C for 3 hours, followed by DNase I treatment to remove DNA template. Finally, synthetic mRNA was purified through LiCl precipitation. The concentration and integrity of mRNA were determined by NanoDrop Spectrophotometer and the Agilent 5200 Fragment Analyzer, respectively.

### Lipid Nanoparticle Preparation and Characterization

mRNA was diluted to 200μg/mL in acetate buffer (pH 4.0). Lipid components (50% ionizable lipid, 38.5% cholesterol, 10% DSPC, and 1.5% PEG-DMG dissolved in ethanol) were mixed with the mRNA solution at a flow rate ratio of 3:1 using a T-mixer device. The resulting LNPs were dialyzed against 2 mM acetate buffer (pH 4.0) using a 100 kDa ultrafiltration membrane to remove ethanol. Finally, the required LNP was obtained after adjusting the pH to 7.5 with Tris buffer.

LNP size and polydispersity index (PDI) were measured using a Zetasizer Pro. Instrument (Malvern). Encapsulation efficiency was determined using the Quant-iT™ RiboGreen Kit (Cat# R11490, Invitrogen). In brief, free mRNA in solution and total mRNA content were measured separately and encapsulation efficiency was calculated as Encapsulation efficiency (%)= (Total mRNA content - Free mRNA content)/Total mRNA content×100%.

### Mammalian Cell Culture

Human embryonic kidney 293T (HEK293T) cells were obtained from the Stem Cell Bank, Chinese Academy of Sciences. Mouse monocyte macrophage RAW 264.7, mouse liver sinusoidal endothelial cells (LSEC), and hepatic stellate cells (HSC) were purchased from Newgainbio Ltd (Wuxi, China). All four cell lines were cultured in Dulbecco’s modified Eagle’s medium (DMEM) (Gibco) supplemented with 10% fetal bovine serum (FBS; Gibco). Primary mouse hepatocytes of C57BL/6 mice were purchased from Milecell Biological Science & Technology Co., Ltd. (Shanghai, China) and were cultured in Maintenance Media. All cells were cultivated at 37℃ with 5%.CO2.

### Animals

C57BL/6 mice (6-8 weeks old, 18-22 g) were purchased from Charles River Laboratory Animal Technology Co., Ltd (Beijing, China) and were cultivated at Ascentage Pharma (Suzhou, China) under standard and pathogen-free conditions (25℃, 50 ± 10% humidity, 12-hour dark/light cycle) with free access to food and water. Animal studies were conducted in accordance with the guidelines of the Chinese Association for Laboratory Animal Sciences and approved by the IACUC of Ascentage Pharma (Approved ethical number: AS-20230621-01, AS-20240313-01, AS-20221228-01).

### Luciferase Assay in Vitro

Cells of HEK293T, RAW 264.7, LSEC, and HSC were seeded into 24-well plates at densities of 2×10⁵, 4×10⁵, 2×10⁵, and 2×10⁵ cells/mL, respectively (500 μL/well), and were grown overnight to the cell density of ∼80%. Next day, cells were transfected with 500 ng/well of mRNA using Lipofectamine™ 3000 Transfection Reagent (Thermo Fisher, Cat. #L3000015). Luciferase assay was performed according to manufacturer’s instruction of Luciferase Assay System kit (Promega, Cat. # E1501) and luminescence signal was measured using Centro Microplate Luminometer (Bethold).

### Luciferase Assay in Vivo

C57BL/6 mice were injected intravenously with *fLuc* mRNA-LNPs at the dose of 1 mg/kg, using 1×PBS as a negative control. At 6 h post-administration, mice were injected intraperitoneally with 150 mg/kg of D-luciferin monosodium salt (YEASEN, Cat. 40901ES01). Mice were subsequently anesthetized in a chamber supplied with 2.5% isoflurane, then placed on the imaging platform while being maintained on 2% isoflurane via a nose cone. The bioluminescence images were taken 10 min after the D-Luciferin injection using IVIS Lumina II (PerkinElmer). After whole-body imaging, mice were sacrificed for organ imaging. The livers and spleens of the same mice were immediately collected and imaged. After all groups were imaged, the average radiance were quantified using IVIS software.

### Statistics

RLU (Relative Luminescence Units) values from in vitro assay or Average Radiance values from in vivo assay of luciferase expression were plotted using GraphPad Prism 10 (GraphPad Software, LLC). Differences between experimental groups in experiments were analyzed by One-way ANOVA multiple comparisons. P-values <0.05 indicate significant differences.

## Acknowledgments

This work was supported by the National Center of Technology Innovation for Biopharmaceuticals NCTIB2023XB02010 (to W.Z.), and CAMS Innovation Fund for Medical Sciences 2021-I2M-1-038 (to S.C.), and 2022-I2M-2-002 (to S.C.).

## Supplementary Figure Legends

**Figure S1.**
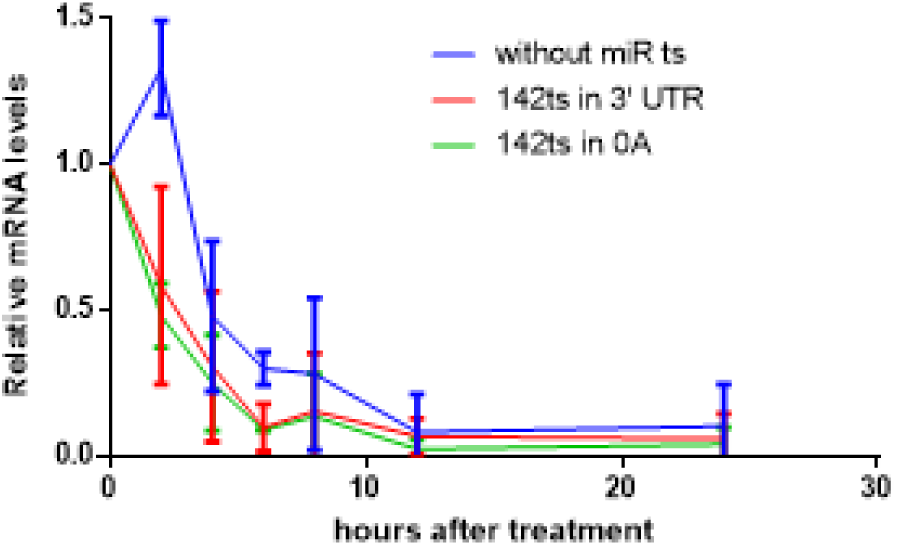
miRNA-142 target sites inserted downstream of 3’ UTR mediate fast mRNA degradation in RAW246.7 macrophages *in vitro*. *fLuc* mRNAs without miR-142 target site insertion (142noMTS) and with the insertion in the 3’ UTR (142MTS-3UTR) or at 0A position of the poly(A) tail (142MTS-0A) were detected by RT-qPCR at different time points after transfecting into RAW246.7 macrophages to determine their degradation rate. GAPDH mRNA was used as internal control.

**Figure S2.**
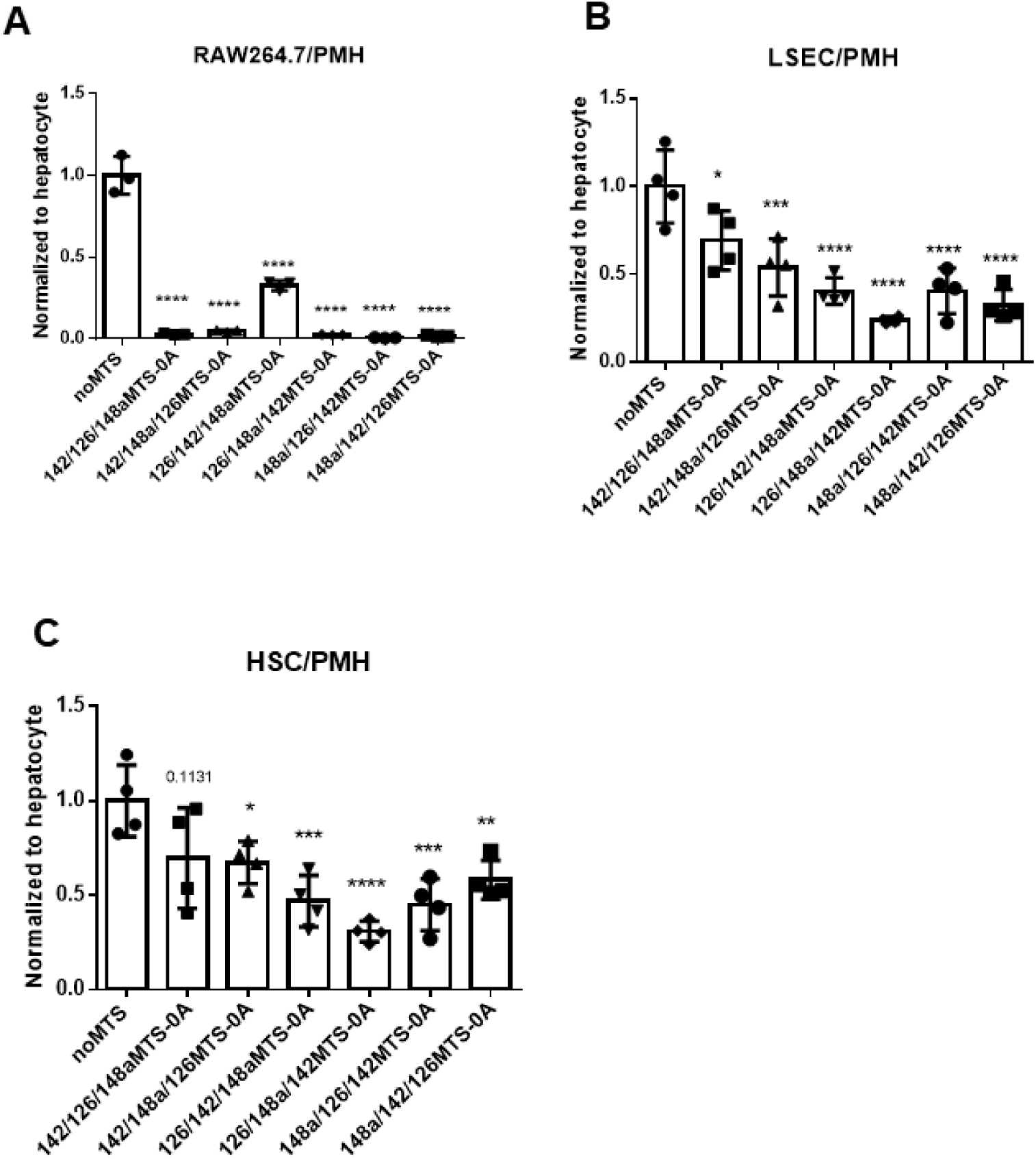
Specific triple miRNA target site insertions (triple-MTS) at the 5’ end of the poly(A) tail efficiently downregulate mRNA expression in three different non-target hepatic cell types. Compared to noMTS control, Triple-MTS insertions for miR-142, miR-126 and miR-148a significantly suppressed luciferase activities in (A) RAW246.7, (B) LSEC, and (C) HSC relative to their activities in primary mouse hepatocytes (PMHs). Data were normalized to noMTS for each graph, with error bars representing SEM (* *p* < 0.05, ** *p* < 0.01, *** *p* < 0.001, **** *p* < 0.0001; one-way ANOVA).

**Supplementary Table 1.**
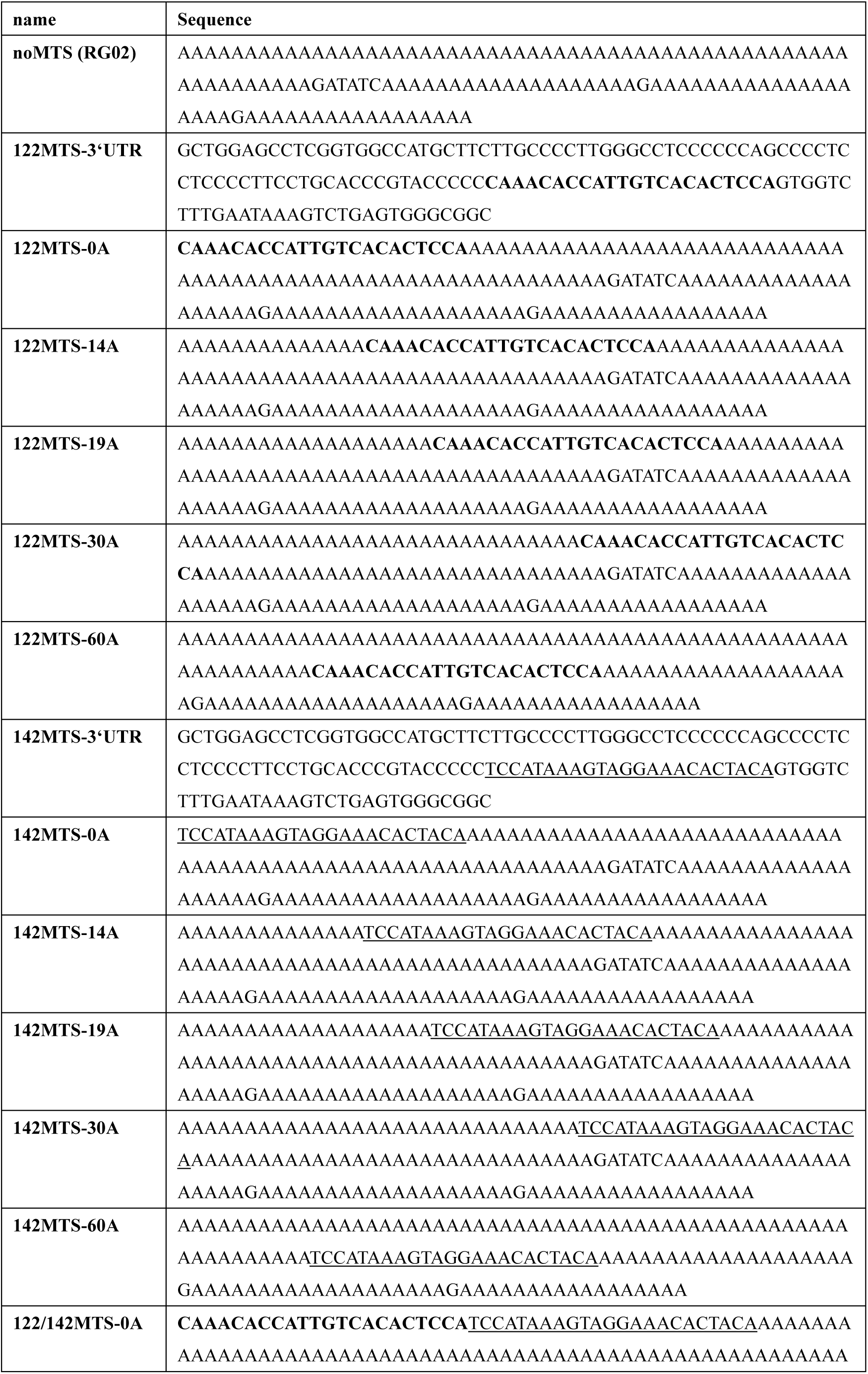
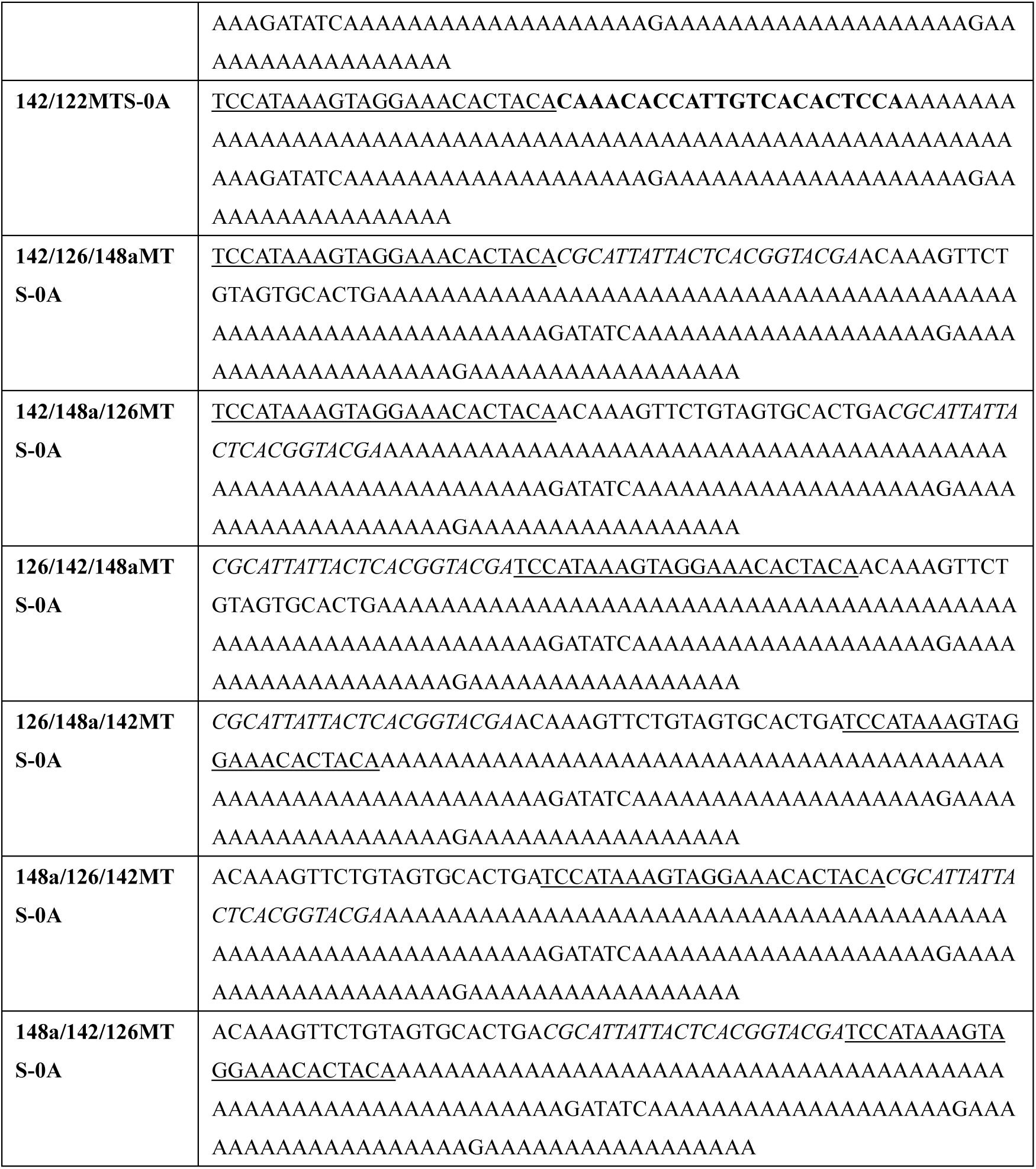
Sequences of poly(A) tails used in this study.

